# EXOC1 regulates cell morphology of spermatogonia and spermatocytes in mice

**DOI:** 10.1101/2020.06.07.139030

**Authors:** Yuki Osawa, Miho Usui, Yumeno Kuba, Hoai Thu Le, Natsuki Mikami, Toshinori Nakagawa, Yoko Daitoku, Kanako Kato, Hossam Hassan Shawki, Yoshihisa Ikeda, Akihiro Kuno, Kento Morimoto, Yoko Tanimoto, Tra Thi Huong Dinh, Kazuya Murata, Ken-ichi Yagami, Masatsugu Ema, Shosei Yoshida, Satoru Takahashi, Seiya Mizuno, Fumihiro Sugiyama

## Abstract

Spermatogenesis requires high regulation of germ cell morphology. The spermatogonia regulates its differentiation state by its own migration. The male germ cells differentiate and mature with the formation of syncytia, failure of forming the appropriate syncytia results in the arrest of spermatogenesis at the spermatocyte stage. However, the detailed molecular mechanisms of male germ cell morphological regulation are unknown. Here, we found that EXOC1 is important for the pseudopod formation of spermatogonia and spermatocyte syncytia in mice. We found that while EXOC1 contributes to the inactivation of Rac1 in the pseudopod formation of spermatogonia, in spermatocyte syncytium formation, EXOC1 and SNAP23 cooperate with STX2. Our results showed that EXOC1 functions in concert with various cell morphology regulators in spermatogenesis. Since EXOC1 is known to bind to several cell morphogenesis factors, this study is expected to be the starting point for the discovery of many morphological regulators of male germ cells.

## Introduction

In mammalian testis, sperms are continuously produced in the seminiferous tubules throughout a male’s life. There are two types of cells in the seminiferous tubule, somatic cells (Sertoli cells) which support spermatogenesis and male germ cells which undergo various processes to become, spermatogonia, spermatocytes, spermatids, and eventually spermatozoa. Structurally, the Sertoli cell tight junction (SCTJ) divides the seminiferous tubule into two compartments, the basal compartment and the luminal compartment (Tsukita, Yamazaki, Katsuno, Tamura, & Tsukita, 2008). Spermatogonia cells are located in the basal compartment and undergo proliferation and differentiation by mitosis. The cells that have differentiated into spermatocytes migrate to the luminal compartment and differentiate into haploid spermatocytes by meiosis (Smith & Braun, 2012). These spermatocytes then undergo dynamic morphological changes to become mature sperms that are released into the lumen of the seminiferous tubule. As this entire differentiation process is cyclical and continuous, long-term stable spermatogenesis is supported by spermatogonial stem cells.

Murine spermatogonia are heterogeneous in their differential state and can be classified as undifferentiated, differentiation-primed, or differentiated (Fayomi & Orwig, 2018) (Suzuki, Sada, Yoshida, & Saga, 2009). Differentiation-primed spermatogonia are largely fated to differentiated permatogonia, but when the number of undifferentiated spermatogonia decreases, these cells dedifferentiate to make up for the loss and function as undifferentiated spermatogonia (Nakagawa, Nabeshima, & Yoshida, 2007). In addition, spermatogonia move within the basal compartment of the seminiferous tubules and regulate the balance between self-renewal and differentiation by competing for fibroblast growth factors (FGFs) that are secreted from deferent niches (Kitadate et al., 2019). Although transplantation studies using in vitro cultured spermatogonial stem cells have reported that Rac1-mediated cell migration is important for spermatogonial stem cell homing (Takashima et al., 2011), the molecules that are involved in spermatogonia migration and their mechanisms have not been clarified (Kanamori, Oikawa, Tanemura, & Hara, 2019).

During spermatogenesis, germ cells undergo an incomplete cytoplasmic division. In most somatic cell types, daughter cells completely separate from each other, but in germ cells, the final separation is inhibited and they differentiate while maintaining intercellular bridges (ICB) (Iwamori et al., 2010). Such structures, called syncytia, are thought to contribute to syncytial differentiation and maturation by the sharing of mRNAs and proteins that are halved during meiosis via ICB (Morales et al., 2002) (Fawcett, Ito, & Slautterback, 1959). TEX14 localizes to germline intercellular bridges and inhibits the final separation of the cytoplasm; in mice lacking this gene, the cytoplasm of male germ cells is completely separated causing spermatogenesis to be halted (Greenbaum et al., 2006). This result indicates that the formation of syncytium is essential for spermatogenesis. Seminolipids, which are sulfated glycolipids, are important for the formation of ICB in spermatocytes. This is evident in mice lacking the *Cgt* & *Cst* genes, required for seminolipids synthesis, as they show an aggregation of spermatocytes (Fujimoto et al., 2000) (Honke et al., 2002). Although the detailed molecular mechanism is unknown, a similar phenotype was observed in mice lacking the *Stx2* gene (Fujiwara et al., 2013), which functions in the transport of seminolipids to the cell membrane. In humans, similar to that of mice, the loss of function of *STX2* results in the failure of spermatogenesis with spermatocyte aggregation (Nakamura et al., 2018).

In this study, we focus on the exocyst complex, a heterodimeric protein complex composed of eight subunits (EXOC1–EXOC8) that is widely conserved from yeast to humans (Koumandou, Dacks, Coulson, & Field, 2007). Exocyst-mediated vesicle trafficking is crucial for various fundamental cellular phenomena such as cell division, morphogenesis, and cell migration. In cytokinesis, the exocyst is localized to ICB (Neto, Balmer, & Gould, 2013) and is required for the transport of factors required for the final separation of the plasma membrane (Kumar et al., 2019). The exocyst has also been implicated in cell migration via the reconstitution of the actin cytoskeleton (Liu et al., 2012) (Parrini et al., 2011).

Although studies using gene-deficient mouse models have shown that the exocyst is important for embryo development (Mizuno et al., 2015) (Friedrich, Hildebrand, & Soriano, 1997) (Fogelgren et al., 2015) (Dickinson et al., 2016), its tissue-specific functions in adults are largely unknown. In *Drosophila* models, the exocyst plays an important role in the formation of both female and male gametes (Giansanti et al., 2015) (Wan et al., 2019) (Murthy & Schwarz, 2004) (Mao et al., 2019). In mice, all exocyst subunits are expressed in male germ cells at each stage of differentiation (Green et al., 2018), and although it is predicted that these subunits may be involved in mammalian spermatogenesis, the function of these subunits is completely unknown. Therefore, in this study, we generated and analyzed a mouse model deficient in *Exoc1*, an exocyst subunit expressed in male germ cells.

## Results

### EXOC1 expression in mouse testis

The expression of *Exoc1* mRNA has been observed in the germ cells of mouse testes at each stage: spermatogonia, spermatocytes, round spermatids, and elongating spermatids (Green et al., 2018). To analyze the expression of EXOC1 at the protein level, *Exoc1*-*PA* knock-in (*Exoc1-PA*) mice were generated using the CRISPR-Cas9 system (Figure 1A) as immunofluorescence using commercially available antibodies for EXOC1 failed to detect any signal. We performed western blot analyses for the PA-tag in the testes of *Exoc1-PA* mice and confirmed a band of the molecular size as expected (Figure 1B). In immunofluorescence, EXOC1-PA was also detected in all testicular cells of *Exoc1-PA* mice (Figure 1C). These data indicate that EXOC1 protein is expressed in male mouse germ cells.

**Figure 1.**
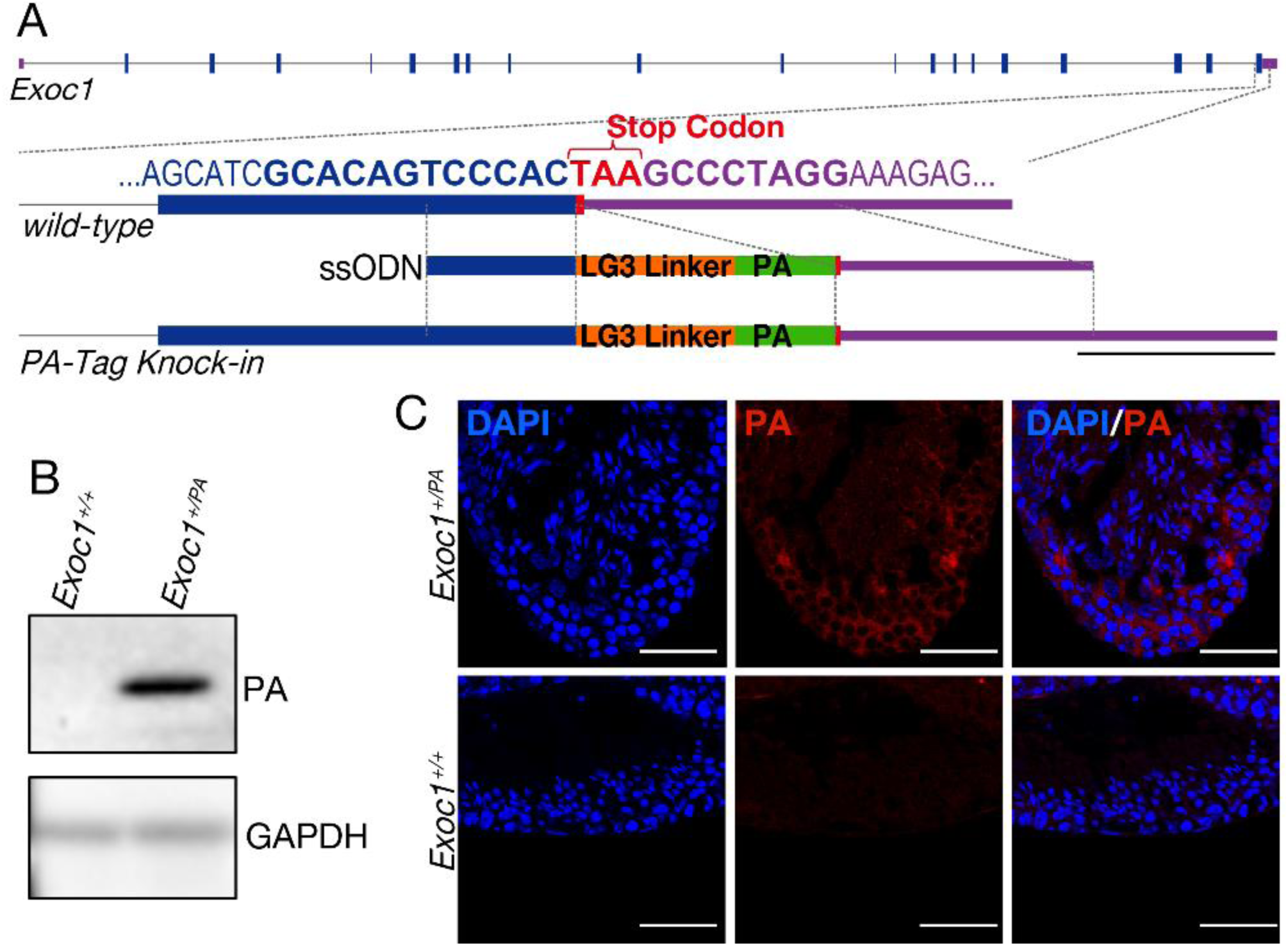
Confirmation of EXOC1 expression in testes using the PA-Tag knock-in mouse. (A) Generation of PA-Tag knock-in mice. The LG3-linker connected PA-tag gene fragment was knocked-in just before the stop codon of *Exoc1* using CRISPR-Cas9. Bold letters represent the CRISPR-Cas9 target sequence. Scale bar: 100 bp. (B) Western-blotting of PA-Tag antibody demonstrated the presence of PA-tagged EXOC1 expression in mouse testis. (C) Immunofluorescence with PA-Tag antibody. EXOC1 is observed in every cell. Scale bars: 50 μm.

### Impaired spermatogenesis in *Exoc1* conditional KO mice

*Exoc1* deficiency results in peri-implantation embryonic lethality in mice (Mizuno et al., 2015). In this study, we analyzed the function of *Exoc1* in spermatogenesis using *Exoc1* conditional knockout (cKO) mice obtained by crossing *Exoc1*-floxed mice (figure supplement 1) with *Nanos3*-*Cre* driver that expresses *Cre* in spermatogonia (Suzuki, Tsuda, Kiso, & Saga, 2008). To investigate the spermatogenic potential of *Exoc1* cKO mice, PNA-lectin staining, to detect acrosome was performed; however, no signal was detected in the *Exoc1* cKO mice (Figure 2A). When cell morphology was observed by H&E staining, although no spermatids and spermatozoa were observed in the testes of *Exoc1* cKO mice, large multinucleated cells were observed in the lumen of seminiferous tubules (Figure 2B). During spermatogenesis, male germ cells form syncytia and are interconnected via stable ICB (Iwamori et al., 2010). Detailed observations by scanning electron microscopy revealed ICB in control mice, while multinucleated cells that lacked ICB structures were observed in *Exoc1* cKO mice (Figure 2C). As the observed multinucleated cells appeared to be aggregates of syncytia, they were named “AGS” for the purpose of this study. To determine at which stage of differentiation the AGS emerge from, immunofluorescence using male germ cell markers for each differentiation stage were used. γH2AX, a spermatocyte marker for the pachytene stage, was observed in the nuclei of AGS (Figure 2D). In contrast, no aggregation was observed in the differentiated spermatogonia, Kit^+^ syncytia (Figure 2E). As the seminiferous tubules are divided into two compartments by the SCTJ, spermatogonia located within the basal compartment migrate to the luminal compartment when they become spermatocytes (Smith & Braun, 2012). In order to investigate the arrangement of AGS in the seminiferous tubules, H&E staining and immunofluorescence for CLDN11, a major component of SCTJ (Morita, Sasaki, Fujimoto, Furuse, & Tsukita, 1999), were performed, and all of the 336 AGS in *Exoc1* cKO mice were observed within the luminal compartment (Figure 2F). These data suggest that the AGS seen in *Exoc1* cKO mice are spermatocytes.

**Figure 2.**
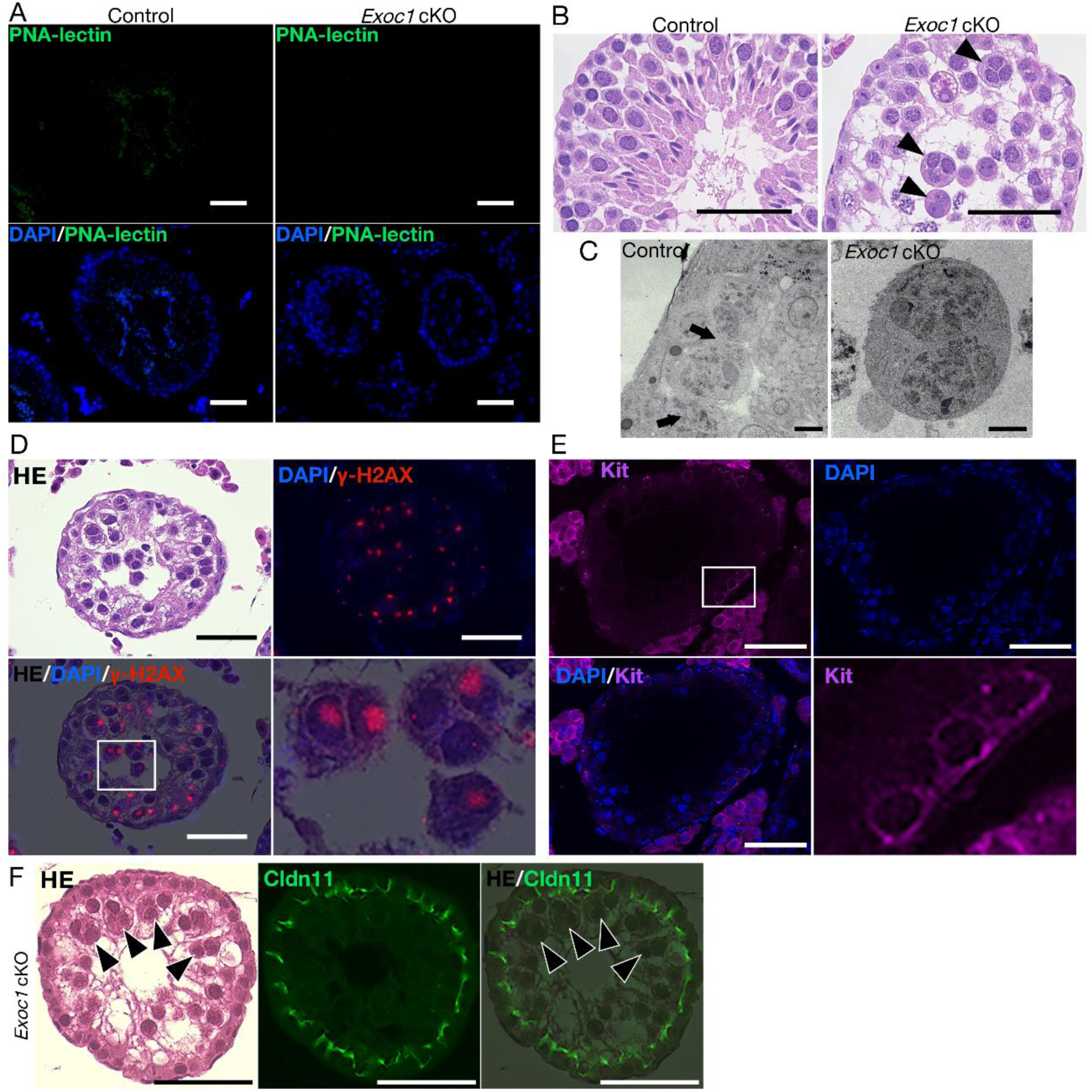
Impaired spermatogenesis in *Exoc1* cKO mice. (A) PNA-lectin staining of the *Exoc1* cKO testis. Signals of PNA-lectin, an acrosomal marker were not observed in the *Exoc1* cKO testis. Scale bars: 50 μm. (B) H&E staining of the *Exoc1* cKO testis. Large and circular cells, appearing to be aggregates of syncytia (AGS), containing multiple nuclei were observed in the lumen of seminiferous tubules (arrowheads). Scale bars: 100 μm. Control: *Exoc1*^*flox/wt*^:: *Nanos3*^*+/Cre*^ mice. (C) SEM observation of the *Exoc1* cKO testis. Intercellular bridges (ICB) (arrows) were found in the syncytia in the control (*Exoc1*^*flox/wt*^:: *Nanos3*^*+/Cre*^) testis. There were no ICB observed in the AGS of *Exoc1* cKO. Scale bars: 5 μm. (D) The serial section overlay image of *Exoc1* cKO testis. There were γ-H2AX (marker of spermatocyte) signals in the AGS nucleus. Scale bars: 50 μm. (E) A representative immunofluorescence image of an *Exoc1* cKO seminiferous tubule. Kit^+^ syncytia, which are observed in differentiating spermatogonia, have ICB. Scale bars: 50 μm. (F) H&E staining and immunofluorescence with CLDN11. CLDN11-positive Sertoli cell tight junction (SCTJ) divides the space between the basal and the luminal compartment, and AGS are present within the luminal compartment (arrowheads). Scale bars: 50 μm.

In the pachytene stage, chromosome synapsis occurs and the synaptonemal complex is formed (Heyting, 1996). To confirm this synapsis, chromosomes of spermatocytes from *Exoc1* cKO mice were spread and immunofluorescence was performed using antibodies against SYCP1 and SYCP3, which are components of the synaptonemal complex. SYCP1 and SYCP3 signals were co-localized in autosomal pairs of the spermatocytes from *Exoc1* cKO mice. These results suggest that meiosis proceeds normally to synapsis in Exoc1-deficient spermatocytes (figure supplement 2).

### EXOC1 regulates ICB formation in cooperation with STX2 and SNAP23

The spermatocytes of *Stx2*^*repro34*^ mice, a null mutant of the *Stx2* gene encoding the SNARE protein, exhibit multinucleated AGS (Fujiwara et al., 2013). Both *Stx2*^*repro34*^ and *Exoc1* cKO mice show similar phenotypes of multinucleated AGS and normal chromosome synapsis (Fujiwara et al., 2013). In addition, in yeast, Sec3 (an ortholog of mouse *Exoc1*) promotes membrane fusion between transport vesicles and the cell membrane by coordinating with Sso2 (an ortholog of mouse *Stx2*) and another SNARE protein, Sec9 (an ortholog of mouse *Snap23*) (Yue et al., 2017). Therefore, we hypothesized that EXOC1 is required for normal syncytia formation in mouse spermatocytes to function in cooperation with STX2 and SNPA23. In yeast, Sec3 directly interacts with Sso2 and promotes the Sso2–Sec9 binary complex formation (Yue et al., 2017). As there were no previous reports regarding the interaction between EXOC1 and STX2 in mice, we analyzed the binding of EXOC1-STX2 and STX2-SNAP23 by the co-immunoprecipitation of HEK293T cells overexpressing EXOC1, STX2, and SNAP23 from mice with FLAG, HA and MYC epitope tags, respectively. Immunoprecipitation of STX2 with anti-HA antibodies in cells overexpressing EXOC1 and STX2 resulted in the detection of EXOC1 bands (Figure 3A). Thus, it was assumed that STX2 interacted with mouse EXOC1 in vitro. Similarly, the binding of STX2 to SNAP23 was also confirmed in cells overexpressing STX2 and SNAP23 and in cells overexpressing these three proteins (Figure 3). These results, consistent with those reported in yeast (Yue et al., 2017), suggested that mouse EXOC1 could bind to STX2 and STX2 to SNAP23.

**Figure 3.**
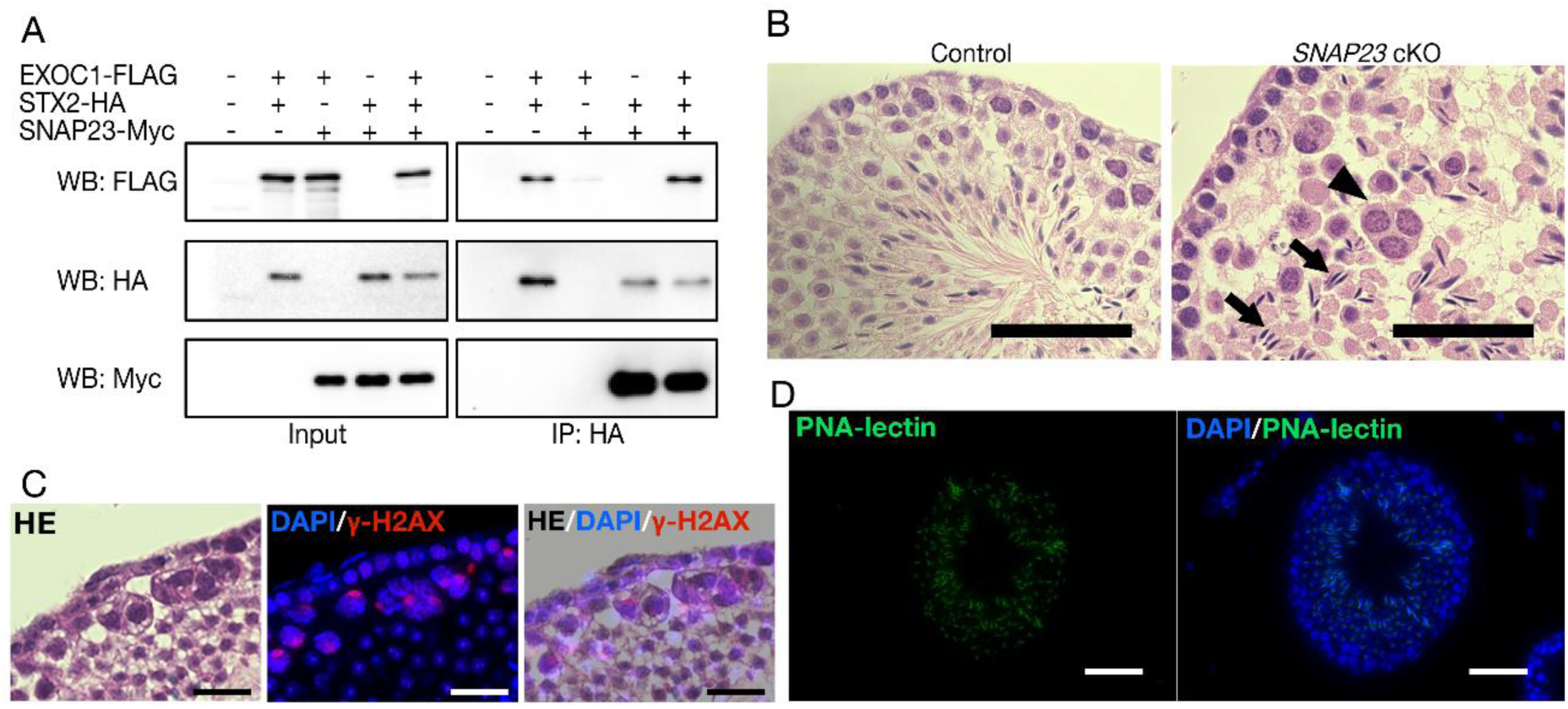
EXOC1 regulates ICB formation in cooperation with STX2 and SNAP23. (A) Co-immunoprecipitation of EXOC1-STX-SNAP23 complex. FLAG-tagged mouse EXOC1, HA-tagged mouse STX2, and Myc-tagged mouse SNAP23 were co-overexpressed in HEK293T cells. Both FLAG-tagged EXOC1 and Myc-tagged SNAP23 were detected by in a HA immunoprecipitated sample. (B) H&E staining of *Snap23* cKO testis. Large and circular cells containing multiple nuclei (arrowheads), appearing to be aggregates of syncytia (AGS) were observed. In contrast with *Exoc1* cKO, every seminiferous tubule had sperms with elongated nuclei (arrows). Scale bars: 50 μm. Control: *Snap23*^*flox/wt*^:: *Nanos3*^*+/Cre*^ mice. (C) The serial section overlay image of *Snap23* cKO testis. Immunofluorescence signals of γ-H2AX were found in AGS. Scale bars: 50 μm. (D) PNA-lectin staining of *Snap23* cKO testis. PNA-lectin that was used to detect the acrosome is observed in the lumen of the seminiferous tubule of *Snap23* cKO testis. Scale bars: 50 μm.

The co-immunoprecipitation described above confirmed that EXOC1, STX2, and SNAP23 interact in mice, and given the hypothesis that EXOC1 functions in cooperation with STX2 and SNAP23 to form normal syncytium of spermatocytes. To test this hypothesis, we conducted a functional analysis of SNAP23. As *Snap23* KO mice have been reported to be embryonically lethal (Suh et al., 2011) and the effect of *Snap23* depletion on adult male germ cells was unknown, we generated *Snap23* cKO mice specifically for male germ cells under the control of *Nanos3* expression (figure supplement 3) and tested for the appearance of spermatocyte AGS. H&E staining and immunofluorescence showed that AGS with γH2AX signals were observed in *Snap23* cKO mice (Figure 3B and C). However, as many spermatocytes did not aggregate in *Snap23* cKO mice, spermatozoa were also observed by H&E and PNA-lectin staining (Figure 3B and D). This suggests that *Snap23* is dispensable for spermatogenesis and that another protein in the SNAP family could be compensating for that function. These results suggest that EXOC1 regulates the formation of the correct syncytium structure in cooperation with STX2 and SNAP23.

### EXOC1 regulates the pseudopod elongation via Rac1 inactivation

As *Nanos3*-Cre mice express CRE from spermatogonia (Suzuki et al., 2008), the *Exoc1* cKO (*Exoc1*^*Flox/Flox*^:: *Nanos3*^*+/Cre*^) mice used in this study were models in which the *Exoc1* gene was deficient from spermatogonia. Therefore, we investigated the effects of *Exoc1* deficiency in spermatogonia. In a study of cultured cells, the exocyst complex was noted to contribute to cell migration by promoting actin assembly (Liu et al., 2012) and GFRα1^+^ undifferentiated spermatogonia were observed to be moving within the basal compartment of the seminiferous tubules (Hara et al., 2014). For these reasons, we carried out detailed morphological observations of GFRα1^+^ spermatogonia using confocal microscopy. The pseudopods of GFRα1^+^ spermatogonia in *Exoc1* cKO mice were significantly shorter than that of control mice (Figure 4A). In the formation of syncytia in spermatocytes, *Exoc1* is likely to function in concert with *Stx2* (Figure 3). In contrast, as there are no reports on the function of *Stx2* in the formation of pseudopods in spermatogonia, it is unclear whether *Exoc1* cooperates with *Stx2* in this phenomenon as well. Therefore, we generated *Stx2* KO mice using the CRISPR-Cas9 system (figure supplement 3A and B) and performed detailed morphological observations of GFRα1^+^ spermatogonia. In the *Stx2* KO mice generated in this study, as in the *Stx2*^*repro34*^ mice (Fujiwara et al., 2013), spermatocytes were observed to aggregate (figure supplement 3C), but no abnormalities in the length of the spermatogonia pseudopod of the *Stx2* KO mice were observed (Figure 4A). In addition, pseudopod length was quantitatively assessed by Sholl analysis: while GFRα1^+^ spermatogonia in *Exoc1* cKO mice had significantly shorter pseudopods than those of the controls (Figure 4B), *Stx2* KO mice did not differ compared to wild-type (Figure 4C). These results suggest that EXOC1 functions in the pseudopod elongation of GFRα1^+^ spermatogonia independently of STX2.

**Figure 4.**
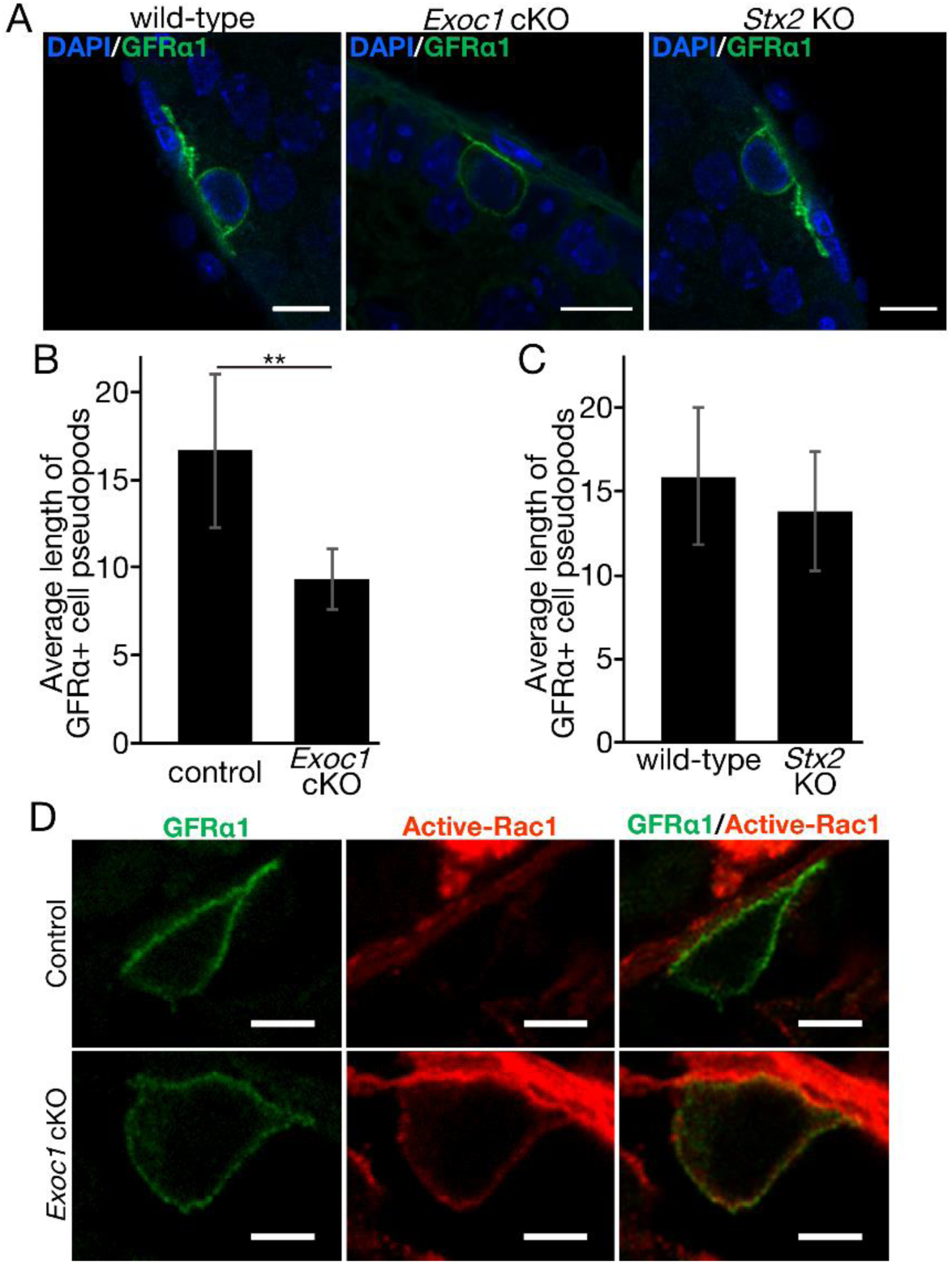
EXOC1 regulates pseudopod elongation via Rac1 inactivation. (A) A representative image of GFRα1^+^ undifferentiated spermatogonia in *Exoc1* cKO and *Stx2* KO. Pseudopod elongation was impaired in *Exoc1* cKO, but not in *Stx2* KO. Scale bars: 10 μm. (B) Pseudopod length quantification. Average length of GFRα1^+^ spermatogonia pseudopods in *Exoc1* cKO was shorter than that of control (n=3, 23∼25 cells in each mouse). **p<0.01, student’s t-test. Control: *Exoc1*^*flox/flox*^ mice. (C) Average length of GFRα1^+^ spermatogonia pseudopods was not statistically significant different between *Stx2* KO and wild type mice. (n=3, 40∼44 cells in each mouse, student’s t-test). (D) Immunofluorescence of active-Rac1 in GFRα1^+^ spermatogonia of *Exoc1* cKO testis. In control mice (*Exoc1*^*flox/wt*^:: *Nanos3*^*+/Cre*^), active-Rac1 signal was lower than the detection limit. Non-polar active-Rac1 signal was detected in *Exoc1* cKO. Scale bars: 5 μm.

We examined the molecular mechanism by which EXOC1 functions in GFRα1^+^ spermatogonia pseudopod elongation. Although the exocyst contributes to pseudopod formation by promoting Arp2/3-mediated actin assembly (Liu et al., 2012), Arp3 is not expressed in spermatogonia (Lie, Chan, Mruk, Lee, & Cheng, 2010). Thus, we focused on the Rho family GTPase Rac1, which regulates pseudopod elongation and cell migration by regulating the reconstruction of the actin cytoskeleton (Raftopoulou & Hall, 2004). The exocyst is required for the transport of SH3BP1, which converts Rac1 from its active to inactive form, and when the exocyst is impaired in cultured human cells, Rac1 is over-activated and pseudopod elongation and cell migration are inhibited (Parrini et al., 2011). Therefore, immunofluorescence for active-Rac1 was performed to examine Rac1 activity in GFRα1^+^ spermatogonia of *Exoc1* cKO mice, and as expected, the fluorescence intensity of active-Rac1 tended to be stronger for the plasma membrane of GFRα1^+^ spermatogonia of *Exoc1* cKO mice compared to control mice (Figure 4D). Therefore, EXOC1 may function in GFRα1^+^ spermatogonia pseudopod elongation by negatively regulating the activity of Rac1.

### Exoc1 functions to regulate the differentiation of spermatogonia

As spermatogonia migration dictates its differentiation state (Kitadate et al., 2019), we counted the number of spermatogonia in two differentiation states GFRα1^+^ cells in the undifferentiated state and RARγ^+^ cells in the differentiation-primed state (Kitadate et al., 2019). Immunofluorescence with antibodies to GFRα1 and RARγ showed a significant increase in undifferentiated GFRα1^+^ cells and a significant decrease in differentiation-primed RARγ^+^ cells in *Exoc1* cKO mice compared to the control mice (Figure 5A and B). In addition, there was no difference in the total number of both cells (Figure 5B). Taken together, these results suggest that *Exoc1* functions to regulate spermatogonia differentiation but not its proliferation.

**Figure 5.**
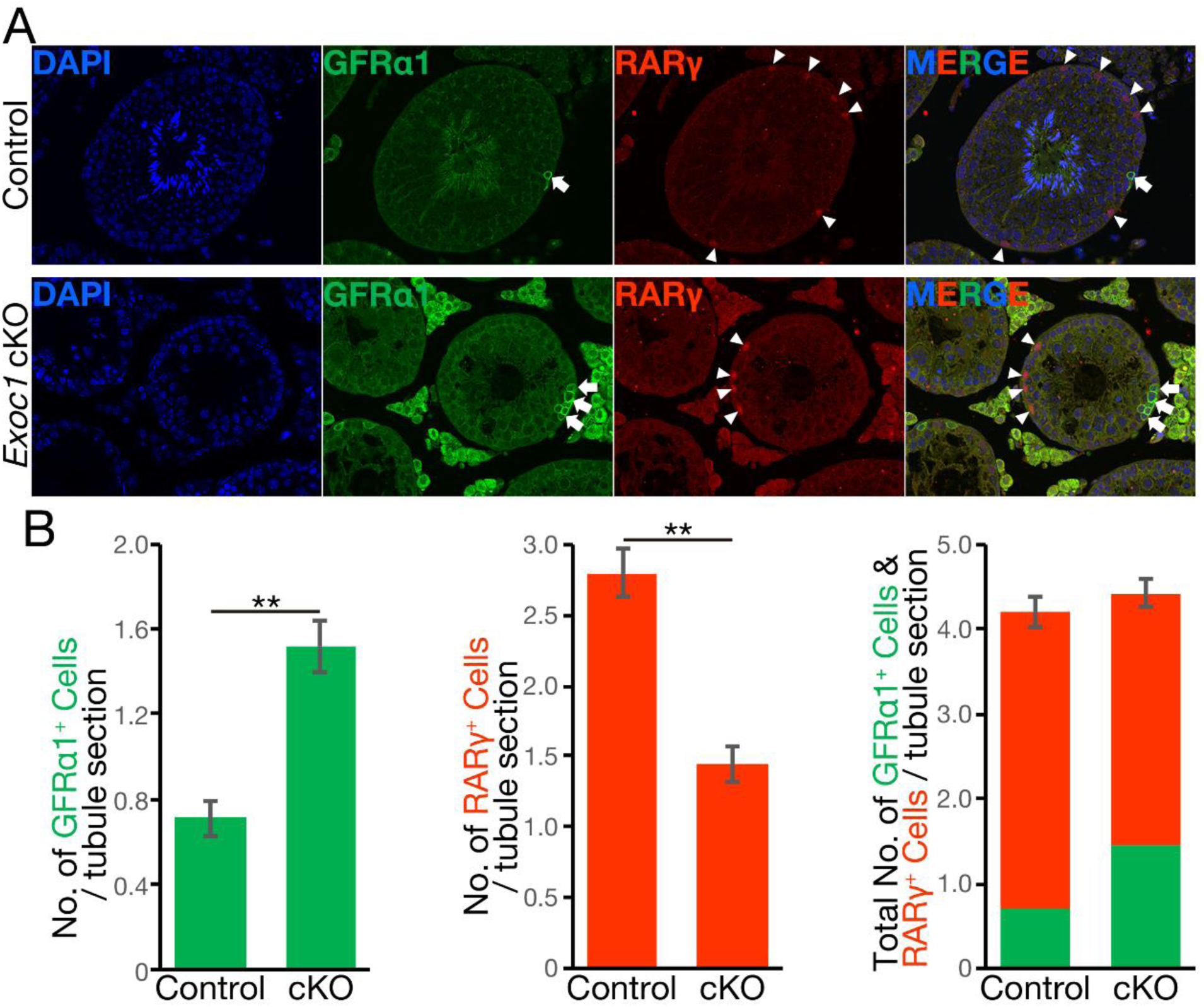
EXOC1 regulates undifferentiated and differentiation-primed spermatogonia density homeostasis. (A) representative immunofluorescence image of *Exoc1* cKO seminiferous tubule. GFRα1^+^ undifferentiated spermatogonia (arrow) density in the cKO was higher than that in control mice. RARγ^+^ differentiation-primed spermatogonia (arrowhead) density decreased in the cKO. Control: *Exoc1*^*flox/flox*^ mice. (B) The number of spermatogonia in a cross section of seminiferous tubule (n=3, 164~188 sections in each mouse). The number of GFRα1^+^ cell per section was significantly increased in the cKO. The number of RARγ^+^ cell in the cKO was significantly smaller than that in control (*Exoc1*^*flox/flox*^) mice. **p<0.01, student’s t-test.

## Discussion

This study is the first to show that EXOC1 plays an essential role in spermatogenesis, in particular the formation of germ cell morphology, in the elongation of the pseudopod in spermatogonia, and in the formation of syncytium in spermatocytes.

*Exoc1* cKO mice showed overactivity of Rac1 and disturbance of differentiation balance in spermatogonia. However, the presence of the next stage of differentiation, spermatocytes, suggests that abnormalities in the spermatogonia are not a direct factor in the failure of spermatogenesis exhibited by the *Exoc1* cKO mice. In contrast, as spermatocytes failed to maintain ICB and aggregated, there were no germ cells observed after spermatid stage. Since chromosome synapsis is suggested to be normal in meiosis, the most significant factor in spermatogenesis failure may be the aggregation of spermatocytes. ICB complex proteins bind to small RNAs, suggesting that ICB may be involved in epigenetic regulation as well as the morphological formation of spermatocytes (Iwamori, Iwamori, Matsumoto, Imai, & Ono, 2020); in *Exoc1* cKO spermatocytes, this epigenetic regulation may be responsible for the arrest of spermatogenesis.

There are three possible reasons for spermatocyte aggregation due to impaired vesicle transport via EXOC1, STX2 and SNAP23; First, the impaired transport of sulfated glycolipids: 90% of the glycolipids in the testes are seminolipids (Ishizuka, 1997), and there are spermatocyte aggregates even in mice lacking the genes required for seminolipid biosynthesis (*Cgt* and *Cst*) (Fujimoto et al., 2000), (Honke et al., 2002). As the localization of seminolipids to the plasma membrane is impaired in spermatocytes of *Stx2*^repro34^ mice (Fujiwara et al., 2013), similarly in *Exoc1* cKO mice, impaired seminolipid transport may have caused for the aggregation of the spermatocytes. Second, insufficient membrane addition: cytokinesis requires the expansion of the surface area of the cell with the formation of cleavage furrow. The exocyst localizes to ICB for local membrane addition (Neto et al., 2013), and mutant *Drosophila*, which lacks *Exoc8* function, shows little expansion of spermatocyte surface area causing the emergence of a defective cytokinetic ring in spermatocytes (Giansanti et al., 2015). This suggests that spermatocyte aggregation in *Exoc1* cKO mice is due to an impaired supply of cell membranes to facilitate cytokinesis. Third, the impaired transport of endosomal sorting complex required for transport (ESCRT) proteins that functions in cell separation: in cytokinesis, cells are separated by ESCRT proteins after the cleavage furrow is squeezed by a contractile ring (Elia, Sougrat, Spurlin, Hurley, & Lippincott-Schwartz, 2011). In cultured human cells, depletion of EXOC3 and EXOC4 impairs the transport of CHMP2B and CHMP4B, subunits of the ESCRT III complex, to ICB and causes multinucleation (Kumar et al., 2019). On the Mouse Genome Informatics database (http://www.informatics.jax.org/), there are ten genes encoding the ESCRT III complex subunit, and two of them (*Chmp2b* and *Chmp4c*) KO mice are fertile (Dickinson et al., 2016). The function of the other eight genes in spermatogenesis has not been analyzed (Coulter et al., 2018), (Lee, Beigneux, Ahmad, Young, & Gao, 2007). Further research will be needed to test these possibilities.

Analysis of the differentiation status of spermatogonia showed an increase in undifferentiated GFRα1^+^ spermatogonia and a decrease in differentiation-primed Rarγ+ spermatogonia in *Exoc1* cKO mice, suggesting that spermatogonia are biased toward undifferentiated maintenance. In addition, the pseudopod of the undifferentiated GFRα1^+^ spermatogonia was shorter in the *Exoc1* cKO mice and the fluorescence intensity of active-Rac1 was stronger. The exocyst is required for the transport of SH3BP1, which converts Rac1 from its active to inactive form, and over-activation of Rac1 inhibits cell migration (Parrini et al., 2011). Therefore, EXOC1 may regulate pseudopod elongation and migration in the undifferentiated spermatogonia by inactivating Rac1 via SH3BP1 transport; since there are no reports of *Sh3bp1* KO mice, it will be necessary to investigate whether pseudopod elongation and migration are impaired when *Sh3bp1* is deficient in undifferentiated spermatogonia. In addition, since undifferentiated spermatogonia move within the basal compartment of the seminiferous tubules and regulate the balance between self-renewal and differentiation by competing with each other for fibroblast growth factors (FGFs) (Kitadate et al., 2019), undifferentiated spermatogonia migration may be important for maintaining the balance between self-renewal and differentiation. The disruption of this balance seen in *Exoc1* cKO mice may have been caused secondary to impaired pseudopod elongation of the cells. In other words, undifferentiated spermatogonia, which normally move within the basal compartment of the seminiferous tubules, replicate themselves near FGFs-producing cells and differentiate into differentiation-primed spermatogonia by leaving FGFs-producing cells via cell migration. In contrast, *Exoc1* deficiency results in insufficient inactivation of Rac1 in undifferentiated spermatogonia and impaired its pseudopod elongation. As a result, undifferentiated spermatogonia that are restricted by migration may remain physically close to FGFs-producing cells and thus repeatedly self-renew, limiting their differentiation into differentiation-primed spermatogonia.

The most serious case of male infertility is azoospermia, which can be divided into two types: obstructive azoospermia, in which the flow of seminiferous tubule fluid is obstructed, and nonobstructive azoospermia, which is due to a problem with the within the testes. In the former case, sperms can be recovered by surgery, but in the latter case, sperms may not be produced at all. A case of spermatogenesis failure with aggregation of spermatocytes due to loss of STX2 function was also reported in humans (Nakamura et al., 2018). The present study suggests that mouse EXOC1 functions in cooperation with mouse STX2 to maintain spermatocyte morphology, proposing that EXOC1 may also be one of the genes responsible for male infertility in humans.

## Materials and methods

### Animals

The mice were maintained in plastic cages under specific pathogen-free conditions in 23.5°C ± 2.5°C and 52.5% ± 12.5% relative humidity under a 14-h light/10-h dark cycle in the Laboratory Animal Resource Center at the University of Tsukuba. Mice had free access to commercial chow (MF diet; Oriental Yeast Co. Ltd) and filtered water. ICR and C57BL/6 mice were purchased from Charles River Laboratories Japan. Nanos3-Cre mice, *Nanos3*^*tm2 (cre)Ysa*^, were kindly gifted by Dr. Saga (Suzuki et al., 2008) through RIKEN BioResource Research Center (RBRC02568). *Exoc1*^*tm1a (EUCOMM)Hmgu*^ and *Snap23*^*tm1a(EUCOMM)Hmgu*^ mice were obtained from the International Knockout Mouse Consortium and the International Mouse Phenotyping Consortium (Skarnes et al., 2011). All male mice used in the experiment were at least 10 weeks old.

### Genome editing in mouse embryos

The *Exoc1* PA Knock-in (KI) mouse were generated by CRISPR-Cas9 based genome editing. The sequence (5′-GCA CAG TCC CAC TAA GCC CT-3′) was selected as the guide RNA (gRNA) target. The gRNA was synthesized and purified by the GeneArt Precision gRNA Synthesis Kit (Thermo Fisher Scientific, Waltham, Massachusetts) and dissolved in Opti-MEM (Thermo Fisher Scientific, Waltham, Massachusetts). In addition, we designed a 200-nt single-stranded DNA oligonucleotide (ssODN) donor; the LG3 linker (Kagoshima et al., 2007) and the PA tag sequence was placed between 55-nt 5′- and 53-nt 3′-homology arms. This ssODN was ordered as Ultramer DNA oligos (Integrated DNA Technologies) and dissolved in Opti-MEM. Two gRNA targets (5′-CAT AAA GTG GTT GCG CTC TT-3′ and 5′-GCA GAT GTG ATG CTC GGC TG-3′) located on intron 4 and 5 of *Stx2*, respectively were selected for producing the *Stx2* KO mouse. These two gRNAs were synthesized, purified, and dissolved in Opti-MEM as mentioned above.

The mixture of gRNA for *Exoc1* (25 ng/μl), ssODN (100 ng/μl), and GeneArt Platinum Cas9 Nuclease (Thermo Fisher Scientific, Waltham, Massachusetts) (100 ng/μl) were electroplated to the zygotes of C57BL/6J mice (Charles River Laboratories Japan, Yokohama, Japan) by using the NEPA 21 electroplater (Nepa Gene Co. Ltd., Ichikawa, Japan) to produce the *Exoc1* PA KI mouse (Sato et al., 2018). The mixture of two gRNAs for *Stx2* (25 ng/μl, each) and GeneArt Platinum Cas9 Nuclease (100 ng/μl) were electroplated to zygotes for producing the *Stx2* KO mouse. After electroporation, 2-cell embryos were transferred into the oviducts of pseudopregnant ICR female and newborns were obtained.

### Electroporation of mice zygotes with *Flpe* mRNA

A *pT7-Flpe-pA* plasmid was constructed from a *T7-NLS hCas9-pA* plasmid, which was kindly gifted by Dr. Mashimo (Yoshimi et al., 2016), through RIKEN BioResource Research Center (RDB13130). *Cas9* cDNA in the *T7-NLS hCas9-pA* was replaced with an *Flpe* cDNA from *pCAG-Flpe*. This *pCAG-Flpe* was kindly gifted by Dr. Cepko (Matsuda & Cepko, 2007) through Addgene (Plasmid #13787). *Flpe* mRNA was transcribed in vitro from *Nhe*I digested *pT7-Flpe-pA* by using mMESSAGE mMACHINE T7 ULTRA Transcription Kit (Thermo Fisher Scientific, Waltham, Massachusetts).

Female C57BL/6 mice (12 weeks old) were superovulated by intraperitoneal administration of 5 units of pregnant mare serum gonadotropin (ASKA Pharmaceutical Co. Ltd., Tokyo, Japan) and 5 units of human chorionic gonadotropin (ASKA Pharmaceutical Co. Ltd., Tokyo, Japan) with a 48-h interval. In vitro fertilization was performed using the wild-type C57BL/6 oocytes and the *Snap23*^*tm1a(EUCOMM)Hmgu*^ sperms according to standard protocols. Five hours later, *Flpe* mRNA (300 ng/μl) was electroplated to the zygotes by using the NEPA 21 electroplater. The poring pulse was set to voltage: 225 V, pulse width: 2 ms, pulse interval: 50 ms, and number of pulses: +4. The transfer pulse was set to voltage: 20 V, pulse width: 50 ms, pulse interval: 50 ms, and number of pulses: ±5 (attenuation rate was set to 40%). A day after electroporation, the developed 2-cell embryos were transferred to pseudopregnant ICR mice.

### Genotyping PCR

Genomic DNA was extracted from < 0.5 mm tails of 3 weeks old mice. PCR was carried out using AmpliTaq Gold 360 Master Mix (Thermo Fisher Scientific, Waltham, Massachusetts) with the appropriate primers (Table S1). For DNA sequencing, PCR products were purified with a FastGene Gel/PCR Extraction Kit (Nippon Genetics, Tokyo, Japan) and sequences were confirmed with a BigDye Terminator v3.1 Cycle Sequencing Kit (Thermo Fisher Scientific, Waltham, Massachusetts), FastGene Dye Terminator Removal Kit (Nippon Genetics, Tokyo, Japan), and 3500xL Genetic Analyzer (Thermo Fisher Scientific, Waltham, Massachusetts).

### Western blot analysis

Protein in testes and cultured cells were extracted using T-PER Tissue protein extraction reagent (Thermo Fisher Scientific, Waltham, Massachusetts), subjected to SDS-PAGE, and transferred to Immobilon-P PVDF membranes (Merck-Millipore, Burlington, Massachusetts). The membranes were blocked in 5% skim milk in Tris-buffered saline overnight at 4°C. The membranes were subsequently incubated with primary antibodies (Table S2) for 1 h at 20-25°C. The membrane was then incubated with HRP-linked secondary antibodies (Table S2) for 1 h at 20-25°C. The blots were developed by chemiluminescence using Luminata Crescendo Western HRP Substrate (Merck-Millipore, Burlington, Massachusetts) and visualized by iBrightCL100 (Thermo Fisher Scientific, Waltham, Massachusetts).

### H&E Staining

Testes, with the tunica albuginea removed, were fixed using 10%-Formaldehyde Neutral Buffer Solution (Nacalai Tesque Inc., Kyoto, Japan) overnight. Fixed testes were then soaked in 70% ethanol and embedded into paraffin blocks. Tissues were sliced at 5 μm using a HM335E microtome (Thermo Fisher Scientific, Waltham, Massachusetts). After deparaffinization, xylene was removed from the sections with 100% ethanol. Moreover, subsequently hydrated with 95% ethanol, 70% ethanol, and deionized distilled water. Hydrated sections were stained with Mayer’s Hematoxylin Solution (FUJIFILM Wako Chemicals, Tokyo, Japan) and 1% Eosin Y Solution (FUJIFILM Wako Chemicals, Tokyo, Japan).

### Electron microscope observation

Testes were fixed in 1% OsO_4_ in 0.1 M phosphate buffer (pH 7.4), dehydrated in ethanol and embedded in epoxy resin poly/Bed 812 (Polysciences Inc., Warrington, Pennsylvania). Subsequently, ultra-thin sections were made at 70–80 nm and stained with uranium acetate and lead citrate. A transmission electron microscope, JEM-1400 (JOEL Ltd., Tokyo, Japan), was used for the observation.

### Immunofluorescence and lectin staining

All immunofluorescence experiments except for active-Rac1, SYCP3, and SYCP1 were performed with paraffin sections. The paraffin sections were prepared similar to that of the H&E staining. After deparaffinization and rehydration, sections were permeabilized with 0.25% TritonX-100 in PBS and autoclaved (121°C, 10 min) with Target Retrieval Solution (Agilent Technologies, Santa Clara, California). Sections were incubated with Blocking One Histo (Nacalai Tesque Inc., Kyoto, Japan) for 15 min at 37°C. Primary antibodies (Table S2) that were diluted with Can Get Signal Immunoreaction Enhancer Solution A (Toyobo Co. Ltd., Osaka, Japan) were applied and slides were incubated for 1 h at 37°C. Alexa Fluor-conjugated secondary antibodies (Table S2) were diluted with Can Get Signal Immunoreaction Enhancer Solution A, and applied for 1 h at 37°C. Prolong gold antifade reagent with DAPI (Thermo Fisher Scientific, Waltham, Massachusetts) was used as a mounting media and for DAPI staining. Active-Rac1 immunofluorescence was performed with frozen sections. To prepare frozen sections, the tunica albuginea was removed from the testes, and the testes were fixed with 4% paraformaldehyde overnight at 4°C. Fixed testes were then soaked in 30% sucrose in PBS overnight at 4°C and embedded in Tissue-Tek O.C.T. Compound (Sakura Finetek, Tokyo, Japan). Tissues were sliced at 14 μm using HM525 NX Cryostat (Thermo Fisher Scientific, Waltham, Massachusetts). Permeabilization, blocking, and antibody reactions were performed similar to that used in immunofluorescence with paraffin sections. Meiotic pachytene chromosome spread and immunofluorescence for SYCP1 and SYCP3 (Table S2) was performed as reported previously (Peters, Plug, van Vugt, & de Boer, 1997). Samples were observed under a BZ-9000 fluorescence microscope (Keyence, Osaka, Japan), a SP8 Confocal Laser Scanning Microscopy (Leica microsystems, Wetzlar, Germany), and an LSM 800 with Airyscan (ZEISS, Oberkochen, Germany). PNA-lectin (Table S2) staining was performed similar to that of immunofluorescence.

### Transfection and Co-IP

Testes RNA was extracted from C57BL/6 mouse using NucleoSpin RNA Plus (Nacalai Tesque Inc., Kyoto, Japan). cDNA was synthesized with PrimeScript RT Master Mix (Takara Bio, Kusatsu, Japan) and full-length cording sequences (CDS) of *Exoc1, Snap23*, and *Stx2* were obtained with PrimeSTAR GXL DNA Polymerase (Takara Bio, Kusatsu, Japan) using the appropriate primers (Table S1). Full-length CDSs were introduced into the *pCDNA3.1* mammalian expression vector with epitope-tag sequences. These vectors were transfected into HEK293T cells with PEI MAX (Polysciences Inc., Warrington, Pennsylvania) according to the manufacturer’s protocol. Co-IP with anti-HA antibody was performed with a HA-tagged Protein PURIFICATION kit (Medical & Biological Laboratories Co., Nagoya, Japan) according to the manufacturer’s protocol.

### Quantification of pseudopod elongation

Immunofluorescence for GFRα1 was performed using thinly sectioned (20 µm) paraffin sections. To include the entire cell, Z-stack photography was performed using a SP8 Confocal Laser Scanning Microscopy (Leica microsystems, Wetzlar, Germany). We applied Sholl analysis (Binley, Ng, Tribble, Song, & Morgan, 2014) to quantify pseudopod elongation, drawing concentric circles around the midpoint of cell body width and the intersection with the basement membrane and measuring the distance of the furthest elongated pseudopod.

### Study approval

All animal experiments were carried out in a humane manner with approval from the Institutional Animal Experiment Committee of the University of Tsukuba in accordance with the Regulations for Animal Experiments of the University of Tsukuba and Fundamental Guidelines for Proper Conduct of Animal Experiments and Related Activities in Academic Research Institutions under the jurisdiction of the Ministry of Education, Culture, Sports, Science, and Technology of Japan.

## Acknowledgements

We would like to thank Tokuko Iwamori for advice on the experimental design for syncytia analyses. We are grateful to Narumi Ogonuki and Atsuo Ogura for their helpful discussions. We would like to thank Yoshihiro Miwa, Hiroyuki Sakuma, and Akio Sekikawa for advice on the experimental design for fluorescence imaging. We are grateful to Aya Ikkyu and Tomoyuki Fujiyama for advice on the experimental design for protein analyses. This work was supported by Scientific Research (B) (17H03566: K.Y. and 19H03142: S.M.) from the Ministry of Education, Culture, Sports, Science, and Technology (MEXT).

## References

Binley, K. E., Ng, W. S., Tribble, J. R., Song, B., & Morgan, J. E. (2014). Sholl analysis: a quantitative comparison of semi-automated methods. J Neurosci Methods, 225, 65–70. doi: 10.1016/j.jneumeth.2014.01.017

Coulter, M. E., Dorobantu, C. M., Lodewijk, G. A., Delalande, F., Cianferani, S., Ganesh, V. S., … Walsh, C. A. (2018). The ESCRT-III Protein CHMP1A Mediates Secretion of Sonic Hedgehog on a Distinctive Subtype of Extracellular Vesicles. Cell Rep, 24(4), 973–986 e978. doi: 10.1016/j.celrep.2018.06.100

Dickinson, M. E., Flenniken, A. M., Ji, X., Teboul, L., Wong, M. D., White, J. K., … Murray, S. A. (2016). High-throughput discovery of novel developmental phenotypes. Nature, 537(7621), 508–514. doi: 10.1038/nature19356

Elia, N., Sougrat, R., Spurlin, T. A., Hurley, J. H., & Lippincott-Schwartz, J. (2011). Dynamics of endosomal sorting complex required for transport (ESCRT) machinery during cytokinesis and its role in abscission. Proc Natl Acad Sci U S A, 108(12), 4846–4851. doi: 10.1073/pnas.1102714108

Fawcett, D. W., Ito, S., & Slautterback, D. (1959). The occurrence of intercellular bridges in groups of cells exhibiting synchronous differentiation. J Biophys Biochem Cytol, 5(3), 453–460. doi: 10.1083/jcb.5.3.453

Fayomi, A. P., & Orwig, K. E. (2018). Spermatogonial stem cells and spermatogenesis in mice, monkeys and men. Stem Cell Res, 29, 207–214. doi: 10.1016/j.scr.2018.04.009

Fogelgren, B., Polgar, N., Lui, V. H., Lee, A. J., Tamashiro, K. K., Napoli, J. A., … Lipschutz, J. H. (2015). Urothelial Defects from Targeted Inactivation of Exocyst Sec10 in Mice Cause Ureteropelvic Junction Obstructions. PLoS One, 10(6), e0129346. doi: 10.1371/journal.pone.0129346

Friedrich, G. A., Hildebrand, J. D., & Soriano, P. (1997). The secretory protein Sec8 is required for paraxial mesoderm formation in the mouse. Dev Biol, 192(2), 364–374. doi: 10.1006/dbio.1997.8727

Fujimoto, H., Tadano-Aritomi, K., Tokumasu, A., Ito, K., Hikita, T., Suzuki, K., & Ishizuka, I. (2000). Requirement of seminolipid in spermatogenesis revealed by UDP-galactose: Ceramide galactosyltransferase-deficient mice. J Biol Chem, 275(30), 22623–22626. doi: 10.1074/jbc.C000200200

Fujiwara, Y., Ogonuki, N., Inoue, K., Ogura, A., Handel, M. A., Noguchi, J., & Kunieda, T. (2013). t-SNARE Syntaxin2 (STX2) is implicated in intracellular transport of sulfoglycolipids during meiotic prophase in mouse spermatogenesis. Biol Reprod, 88(6), 141. doi: 10.1095/biolreprod.112.107110

Giansanti, M. G., Vanderleest, T. E., Jewett, C. E., Sechi, S., Frappaolo, A., Fabian, L., … Blankenship, J. T. (2015). Exocyst-Dependent Membrane Addition Is Required for Anaphase Cell Elongation and Cytokinesis in Drosophila. PLoS Genet, 11(11), e1005632. doi: 10.1371/journal.pgen.1005632

Green, C. D., Ma, Q., Manske, G. L., Shami, A. N., Zheng, X., Marini, S., … Hammoud, S. S. (2018). A Comprehensive Roadmap of Murine Spermatogenesis Defined by Single-Cell RNA-Seq. Dev Cell, 46(5), 651–667 e610. doi: 10.1016/j.devcel.2018.07.025

Greenbaum, M. P., Yan, W., Wu, M. H., Lin, Y. N., Agno, J. E., Sharma, M., … Matzuk, M. M. (2006). TEX14 is essential for intercellular bridges and fertility in male mice. Proc Natl Acad Sci U S A, 103(13), 4982–4987. doi: 10.1073/pnas.0505123103

Hara, K., Nakagawa, T., Enomoto, H., Suzuki, M., Yamamoto, M., Simons, B. D., & Yoshida, S. (2014). Mouse spermatogenic stem cells continually interconvert between equipotent singly isolated and syncytial states. Cell Stem Cell, 14(5), 658–672. doi: 10.1016/j.stem.2014.01.019

Heyting, C. (1996). Synaptonemal complexes: structure and function. Curr Opin Cell Biol, 8(3), 389–396. doi: 10.1016/s0955-0674(96)80015-9

Honke, K., Hirahara, Y., Dupree, J., Suzuki, K., Popko, B., Fukushima, K., … Taniguchi, N. (2002). Paranodal junction formation and spermatogenesis require sulfoglycolipids. Proc Natl Acad Sci U S A, 99(7), 4227–4232. doi: 10.1073/pnas.032068299

Ishizuka, I. (1997). Chemistry and functional distribution of sulfoglycolipids. Prog Lipid Res, 36(4), 245–319. doi: 10.1016/s0163-7827(97)00011-8

Iwamori, T., Iwamori, N., Ma, L., Edson, M. A., Greenbaum, M. P., & Matzuk, M. M. (2010). TEX14 interacts with CEP55 to block cell abscission. Mol Cell Biol, 30(9), 2280–2292. doi: 10.1128/MCB.01392-09

Iwamori, T., Iwamori, N., Matsumoto, M., Imai, H., & Ono, E. (2020). Novel localizations and interactions of intercellular bridge proteins revealed by proteomic profiling. Biol Reprod. doi: 10.1093/biolre/ioaa017

Kagoshima, H., Nimmo, R., Saad, N., Tanaka, J., Miwa, Y., Mitani, S., … Woollard, A. (2007). The C. elegans CBFbeta homologue BRO-1 interacts with the Runx factor, RNT-1, to promote stem cell proliferation and self-renewal. Development, 134(21), 3905–3915. doi: 10.1242/dev.008276

Kanamori, M., Oikawa, K., Tanemura, K., & Hara, K. (2019). Mammalian germ cell migration during development, growth, and homeostasis. Reprod Med Biol, 18(3), 247–255. doi: 10.1002/rmb2.12283

Kitadate, Y., Jorg, D. J., Tokue, M., Maruyama, A., Ichikawa, R., Tsuchiya, S., … Yoshida, S. (2019). Competition for Mitogens Regulates Spermatogenic Stem Cell Homeostasis in an Open Niche. Cell Stem Cell, 24(1), 79–92 e76. doi: 10.1016/j.stem.2018.11.013

Koumandou, V. L., Dacks, J. B., Coulson, R. M., & Field, M. C. (2007). Control systems for membrane fusion in the ancestral eukaryote; evolution of tethering complexes and SM proteins. BMC Evol Biol, 7, 29. doi: 10.1186/1471-2148-7-29

Kumar, H., Pushpa, K., Kumari, A., Verma, K., Pergu, R., & Mylavarapu, S. V. S. (2019). The exocyst complex and Rab5 are required for abscission by localizing ESCRT III subunits to the cytokinetic bridge. J Cell Sci, 132(14). doi: 10.1242/jcs.226001

Lee, J. A., Beigneux, A., Ahmad, S. T., Young, S. G., & Gao, F. B. (2007). ESCRT-III dysfunction causes autophagosome accumulation and neurodegeneration. Curr Biol, 17(18), 1561–1567. doi: 10.1016/j.cub.2007.07.029

Lie, P. P., Chan, A. Y., Mruk, D. D., Lee, W. M., & Cheng, C. Y. (2010). Restricted Arp3 expression in the testis prevents blood-testis barrier disruption during junction restructuring at spermatogenesis. Proc Natl Acad Sci U S A, 107(25), 11411–11416. doi: 10.1073/pnas.1001823107

Liu, J., Zhao, Y., Sun, Y., He, B., Yang, C., Svitkina, T., … Guo, W. (2012). Exo70 stimulates the Arp2/3 complex for lamellipodia formation and directional cell migration. Curr Biol, 22(16), 1510–1515. doi: 10.1016/j.cub.2012.05.055

Mao, Y., Tu, R., Huang, Y., Mao, D., Yang, Z., Lau, P. K., … Xie, T. (2019). The exocyst functions in niche cells to promote germline stem cell differentiation by directly controlling EGFR membrane trafficking. Development, 146(13). doi: 10.1242/dev.174615

Matsuda, T., & Cepko, C. L. (2007). Controlled expression of transgenes introduced by in vivo electroporation. Proc Natl Acad Sci U S A, 104(3), 1027–1032. doi: 10.1073/pnas.0610155104

Mizuno, S., Takami, K., Daitoku, Y., Tanimoto, Y., Dinh, T. T., Mizuno-Iijima, S., … Yagami, K. (2015). Peri-implantation lethality in mice carrying megabase-scale deletion on 5qc3.3 is caused by Exoc1 null mutation. Sci Rep, 5, 13632. doi: 10.1038/srep13632

Morales, C. R., Lefrancois, S., Chennathukuzhi, V., El-Alfy, M., Wu, X., Yang, J., … Hecht, N. B. (2002). A TB-RBP and Ter ATPase complex accompanies specific mRNAs from nuclei through the nuclear pores and into intercellular bridges in mouse male germ cells. Dev Biol, 246(2), 480–494. doi: 10.1006/dbio.2002.0679

Morita, K., Sasaki, H., Fujimoto, K., Furuse, M., & Tsukita, S. (1999). Claudin-11/OSP-based tight junctions of myelin sheaths in brain and Sertoli cells in testis. J Cell Biol, 145(3), 579–588. doi: 10.1083/jcb.145.3.579

Murthy, M., & Schwarz, T. L. (2004). The exocyst component Sec5 is required for membrane traffic and polarity in the Drosophila ovary. Development, 131(2), 377–388. doi: 10.1242/dev.00931

Nakagawa, T., Nabeshima, Y., & Yoshida, S. (2007). Functional identification of the actual and potential stem cell compartments in mouse spermatogenesis. Dev Cell, 12(2), 195–206. doi: 10.1016/j.devcel.2007.01.002

Nakamura, S., Kobori, Y., Ueda, Y., Tanaka, Y., Ishikawa, H., Yoshida, A., … Fukami, M. (2018). STX2 is a causative gene for nonobstructive azoospermia. Hum Mutat, 39(6), 830–833. doi: 10.1002/humu.23423

Neto, H., Balmer, G., & Gould, G. (2013). Exocyst proteins in cytokinesis: Regulation by Rab11. Commun Integr Biol, 6(6), e27635. doi: 10.4161/cib.27635

Parrini, M. C., Sadou-Dubourgnoux, A., Aoki, K., Kunida, K., Biondini, M., Hatzoglou, A., … Camonis, J. (2011). SH3BP1, an exocyst-associated RhoGAP, inactivates Rac1 at the front to drive cell motility. Mol Cell, 42(5), 650–661. doi: 10.1016/j.molcel.2011.03.032

Peters, A. H., Plug, A. W., van Vugt, M. J., & de Boer, P. (1997). A drying-down technique for the spreading of mammalian meiocytes from the male and female germline. Chromosome Res, 5(1), 66–68. doi: 10.1023/a:1018445520117

Raftopoulou, M., & Hall, A. (2004). Cell migration: Rho GTPases lead the way. Dev Biol, 265(1), 23–32. doi: 10.1016/j.ydbio.2003.06.003

Sato, Y., Tsukaguchi, H., Morita, H., Higasa, K., Tran, M. T. N., Hamada, M., … Yoshimura, A. (2018). A mutation in transcription factor MAFB causes Focal Segmental Glomerulosclerosis with Duane Retraction Syndrome. Kidney Int, 94(2), 396–407. doi: 10.1016/j.kint.2018.02.025

Skarnes, W. C., Rosen, B., West, A. P., Koutsourakis, M., Bushell, W., Iyer, V., … Bradley, A. (2011). A conditional knockout resource for the genome-wide study of mouse gene function. Nature, 474(7351), 337–342. doi: 10.1038/nature10163

Smith, B. E., & Braun, R. E. (2012). Germ cell migration across Sertoli cell tight junctions. Science, 338(6108), 798–802. doi: 10.1126/science.1219969

Suh, Y. H., Yoshimoto-Furusawa, A., Weih, K. A., Tessarollo, L., Roche, K. W., Mackem, S., & Roche, P. A. (2011). Deletion of SNAP-23 results in pre-implantation embryonic lethality in mice. PLoS One, 6(3), e18444. doi: 10.1371/journal.pone.0018444

Suzuki, H., Sada, A., Yoshida, S., & Saga, Y. (2009). The heterogeneity of spermatogonia is revealed by their topology and expression of marker proteins including the germ cell-specific proteins Nanos2 and Nanos3. Dev Biol, 336(2), 222–231. doi: 10.1016/j.ydbio.2009.10.002

Suzuki, H., Tsuda, M., Kiso, M., & Saga, Y. (2008). Nanos3 maintains the germ cell lineage in the mouse by suppressing both Bax-dependent and -independent apoptotic pathways. Dev Biol, 318(1), 133–142. doi: 10.1016/j.ydbio.2008.03.020

Takashima, S., Kanatsu-Shinohara, M., Tanaka, T., Takehashi, M., Morimoto, H., & Shinohara, T. (2011). Rac mediates mouse spermatogonial stem cell homing to germline niches by regulating transmigration through the blood-testis barrier. Cell Stem Cell, 9(5), 463–475. doi: 10.1016/j.stem.2011.08.011

Tsukita, S., Yamazaki, Y., Katsuno, T., Tamura, A., & Tsukita, S. (2008). Tight junction-based epithelial microenvironment and cell proliferation. Oncogene, 27(55), 6930–6938. doi: 10.1038/onc.2008.344

Wan, P., Zheng, S., Chen, L., Wang, D., Liao, T., Yan, X., & Wang, X. (2019). The Exocyst Component Sec3 Controls Egg Chamber Development Through Notch During Drosophila Oogenesis. Front Physiol, 10, 345. doi: 10.3389/fphys.2019.00345

Yoshimi, K., Kunihiro, Y., Kaneko, T., Nagahora, H., Voigt, B., & Mashimo, T. (2016). ssODN-mediated knock-in with CRISPR-Cas for large genomic regions in zygotes. Nat Commun, 7, 10431. doi: 10.1038/ncomms10431

Yue, P., Zhang, Y., Mei, K., Wang, S., Lesigang, J., Zhu, Y., … Guo, W. (2017). Sec3 promotes the initial binary t-SNARE complex assembly and membrane fusion. Nat Commun, 8, 14236. doi: 10.1038/ncomms14236

