## Supplemental Figures for "EXOC1 regulates cell morphology of spermatogonia and spermatocytes in mice"

**Title**

**Abstract**

Spermatogenesis requires high regulation of germ cell morphology. The spermatogonia regulates its differentiation state by its own migration. The male germ cells differentiate and mature with the formation of syncytia, failure of forming the appropriate syncytia results in the arrest of spermatogenesis at the spermatocyte stage. However, the detailed molecular mechanisms of male germ cell morphological regulation are unknown. Here, we found that EXOC1 is important for the pseudopod formation of spermatogonia and spermatocyte syncytia in mice. We found that while EXOC1 contributes to the inactivation of Rac1 in the pseudopod formation of spermatogonia, in spermatocyte syncytium formation, EXOC1 and SNAP23 cooperate with STX2. Our results showed that EXOC1 functions in concert with various cell morphology regulators in spermatogenesis. Since EXOC1 is known to bind to several cell morphogenesis factors, this study is expected to be the starting point for the discovery of many morphological regulators of male germ cells.

### Supplemental Information

Supplemental Table 1. Primers for genotyping

| Experiment | Primer Name | Sequence |
|---|---|---|
| *Exoc1*$^{PA}$ genotyping | Exoc1 3F screening F | ATGGAGTCATCCCCTCACAG |
| | Exoc1 3F screening R | GTCCCTGGAAAGTGCTGAAG |
| *Exoc1*$^{flox}$ genotyping<br>*Exoc1*$^{cKO}$ genotyping | Exoc1-cKO genotyping n3-1 | GGAAAACCCGGAATTTGATT |
| | Exoc1-cKO genotyping n3-2 | CCTTGCACACCAATGAGCTA |
| | Exoc1-cKO genotyping n3-3 | CCGGATTGATGGTAGTGGTC |
| *Snap23*$^{flox}$ genotyping | Snap23-Genotyping-1 | TGCAGGAATCAAGACCATCA |
| | Snap23-Genotyping-2 | CAGCATGTGAGGGTTGCTTA |
| *Snap23*$^{cKO}$ genotyping | Snap23-Genotyping-1 | TGCAGGAATCAAGACCATCA |
| | Snap23-Genotyping-6 | GCCTTTGAGTGACAAGCTCA |
| *Stx2*$^{KO}$ genotyping | Stx2 QC primer Long F | CAGGTGTTTGAGAAAGTGGGGATAGCTT |
| | Stx2 QC primer Long R | TAAAACTGACGACTAAGAAGCCCCAACC |
| | Stx2 QC primer Short F | AACACGTAAGGCCCATGTTC |
| | Stx2 QC primer Short R | AACCTGGGTCCTTCGTATCC |
| Construction of *Exoc1* expression vector | Exoc1 full length RT-F | ATGACAGCAATCAAGCATGCGCTGCAGAGA |
| | Exoc1 full length RT-R | GTGGGACTGTGCGATGCTGGAGCAATA |
| Construction of *Snap23* expression vector | Snap23 full length RT-F | ATGGATAATCTGTCCCCAGAGGA |
| | Snap23 full length RT-F | ACTATCAATGAGTTTCTTTGCTCTTGTA |
| Construction of *Stx2* expression vector | Stx2 full length RT-F | ATGCGGGACCGGCTGCCC |
| | Stx2 full length RT-F | GACAATGCTGTTGCGAGAATAATTCCA |

Supplemental Table 2. Antibodies

| Experiment | Antibodies | Source | Identifier |
|---|---|---|---|
| Western Blotting | Rat monoclonal anti-PA-tag (used at 1:1000) | Wako Pure Chemical Industries | Cat#016-25861 |
| Western Blotting | Rabbit polyclonal anti-GAPDH (used at 1:1000) | Santa Cruz Biotechnology | Cat#sc25778 |
| Western Blotting | Rabbit polyclonal anti-FLAG (used at 1:1000) | Sigma-Adrich | Cat#F7425 |
| Western Blotting | Rabbit polyclonal anti-HA-tag (used at 1:1000) | MEDICAL & BIOLOGICAL LABORATORIES | Cat#561 |
| Western Blotting | Mouse monoclonal anti-Myc (used at 1:1000) | MEDICAL & BIOLOGICAL LABORATORIES | Cat#M192-3 |
| Western Blotting | Goat anti-Rat IgG, HRP-linked (used at 1:2500) | GE Healthcare | Cat#NA935V |
| Western Blotting | Donkey anti-Rabbit IgG, HRP-linked (used at 1:2500) | GE Healthcare | Cat#NA934V |
| Western Blotting | Sheep anti-Mouse IgG, HRP-linked (used at 1:2500) | GE Healthcare | Cat#NA931V |
| Immunofluorescence | Rat monoclonal anti-PA-tag (used at 1:1000) | FUJIFILM Wako Chemicals | Cat#016-25861 |
| Immunofluorescence | Mouse monoclonal anti-γH2AX (used at 1:100) | Merck-Millipore | Cat#05-636 |
| Immunofluorescence | Rabbit polyclonal anti-SYCP1 (used at 1:50) | Novus Biological | Cat#NB300-299 |
| Immunofluorescence | Mouse monoclonal anti-SYCP3 (used at 1:50) | Santa Cruz Biotechnology | Cat#sc-74569 |
| Immunofluorescence | Goat polyclonal anti-GFRα1 (used at 1:400) | R&D systems | Cat#AF560 |
| Immunofluorescence | Rabbit monoclonal anti-RARγ1 (used at 1:200) | Cell Signaling Technology | Cat#8965S |
| Immunofluorescence | Mouse monoclonal anti-active rac1 (used at 1:1000) | NewEast Biosciences | Cat#26903 |
| Immunofluorescence | Rabbit polyclonal anti-Exoc1 (used at 1:50) | Proteintech | Cat#11690-1-AP |
| Immunofluorescence | Rabbit polyclonal anti-Exoc1 (used at 1:50) | Atlas Antibodies | Cat#HPA037706 |
| Immunofluorescence | Chicken anti-Goat IgG, Alexa Fluor 488 (used at 1:200) | Thermo Fisher Scientific | Cat#A21467 |
| Immunofluorescence | Donkey anti-Rat IgG, Alexa Fluor 555 (used at 1:1000) | Abcam | Cat#ab150154 |
| Immunofluorescence | Goat anti-Mouse IgG, Alexa Fluor 555 (used at 1:200) | Thermo Fisher Scientific | Cat#A28180 |
| Immunofluorescence | Donkey anti-Mouse IgG, Alexa Fluor 555 (used at 1:200) | Thermo Fisher Scientific | Cat#A31570 |
| Immunofluorescence | Goat anti-Rabbit IgG, Alexa Fluor 647 (used at 1:200) | Thermo Fisher Scientific | Cat#A27040 |
| Lectin staining | Lectin from Arachis hypogaea, FITC (used at 1:100) | Sigma-Adrich | Cat#L7381 |

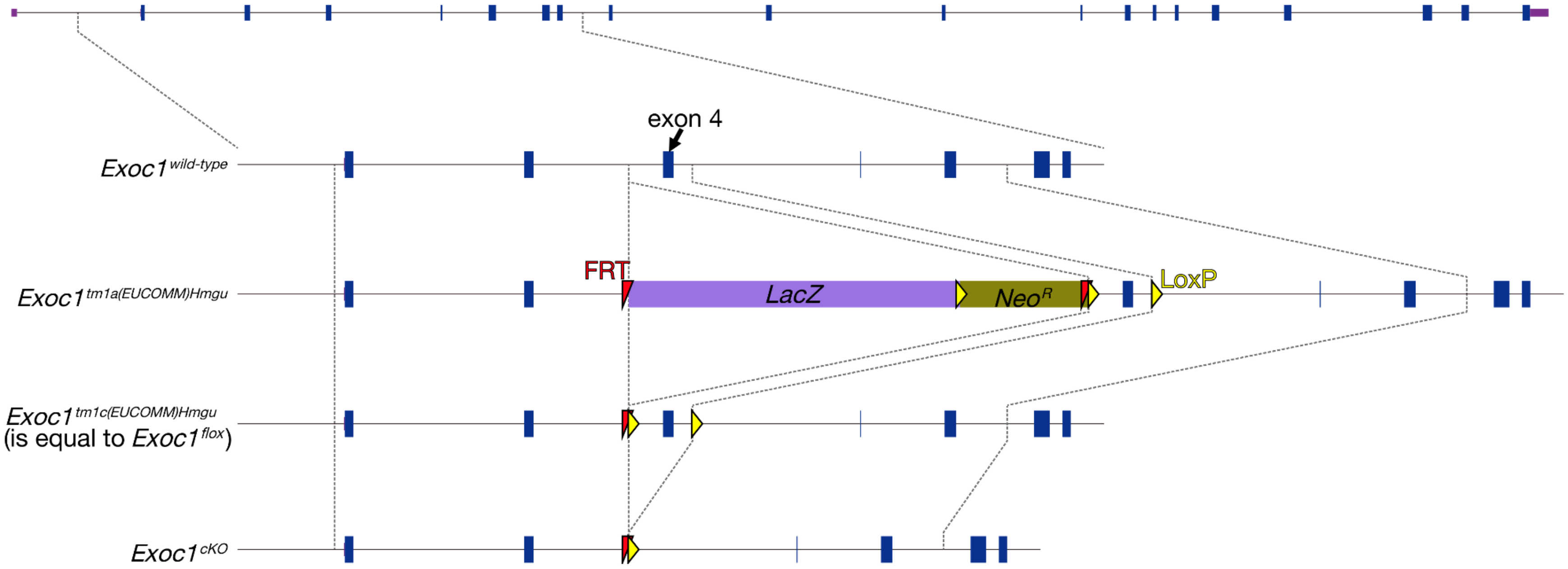


figure supplement 1. The production of *Exoc1* flox mice.

The *Exoc1* flox strain was from the *Exoc1*$^{tm1a(EUCOMM)Hmgu}$ mouse. The *Exoc1*$^{tm1a(EUCOMM)Hmgu}$ allele was changed to the *Exoc1*$^{tm1c(EUCOMM)Hmgu}$ (is equal to *Exoc1*$^{flox}$) allele by mating with the Flpe expression mouse (B6;SJL-Tg(ACTFLPe)9205Dym/J) (Muranishi et al., 2011). Exon 4 of *Exoc1* was floxed in this allele. Arrows indicate primers for detecting the flox allele.

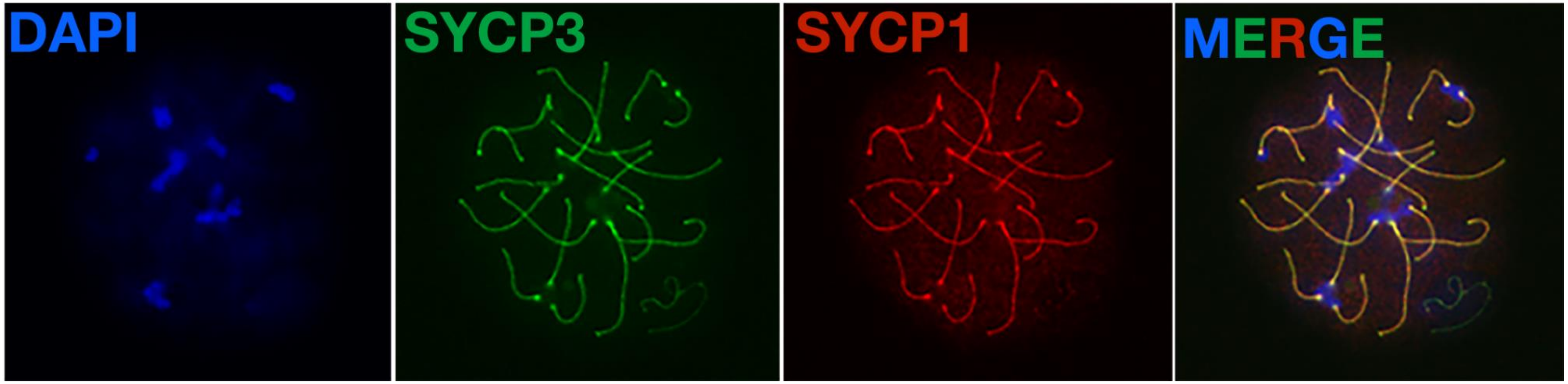


figure supplement 2. Normal meiotic chromosome synapsis in *Exoc1* cKO mice.

Meiotic chromosome synapsis was confirmed by immunofluorescence. In pachytene stage, both the homologous pairing (marked by Sycp3) and synaptonemal complex (marked by Sycp1) were normal in *Exoc1* cKO autosomal chromosomes (n=2).

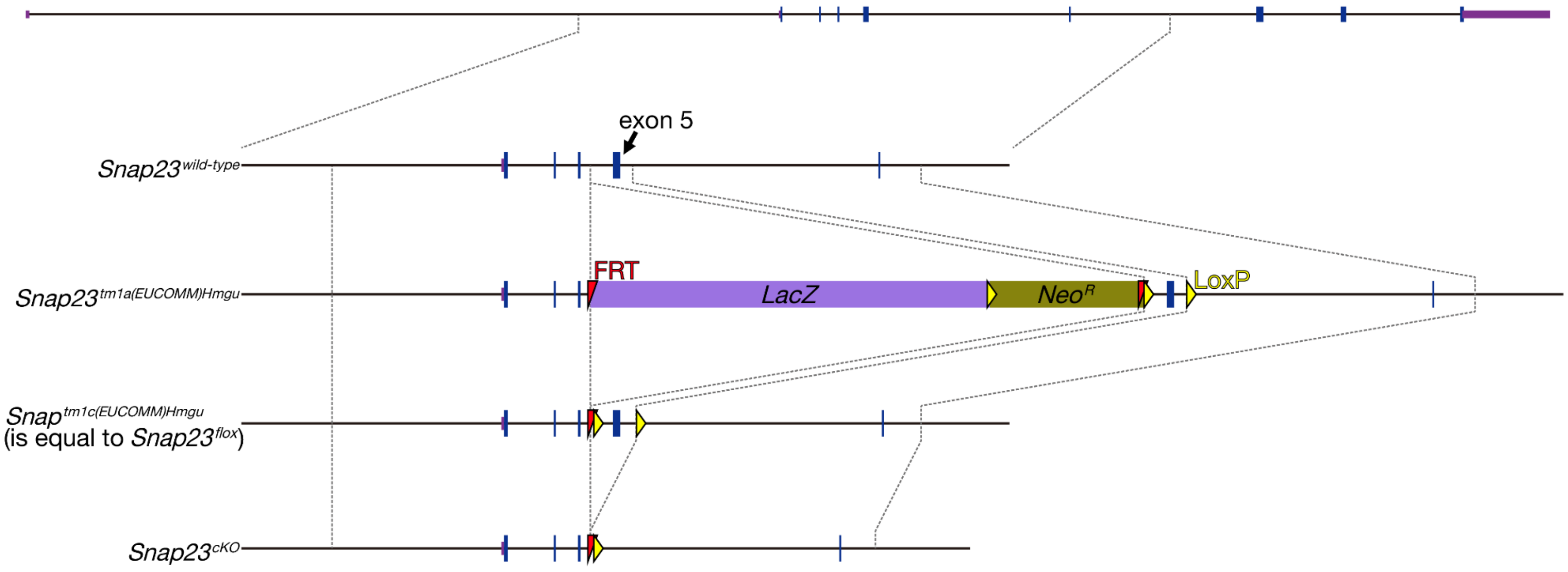


figure supplement 3. The production of *Snap23* flox mice.
*Snap23* flox mice were from the *Snap23*$^{tm1a(EUCOMM)Wtsi}$ frozen sperms. *Snap23*$^{tm1a(EUCOMM)Wtsi}$ frozen sperms were thawed and fertilized to wild-type oocyte in vitro. *Flpe* mRNA was electroplated into the fertilized embryos. *Snap23*$^{tm1a(EUCOMM)Wtsi}$ was changed to *Snap23*$^{tm1c(EUCOMM)Wtsi}$ (equal to *Snap23* flox) allele in every newborn. Exon 5 of *Snap23* was floxed in this allele. Arrows indicate the primers for the detection of flox allele.

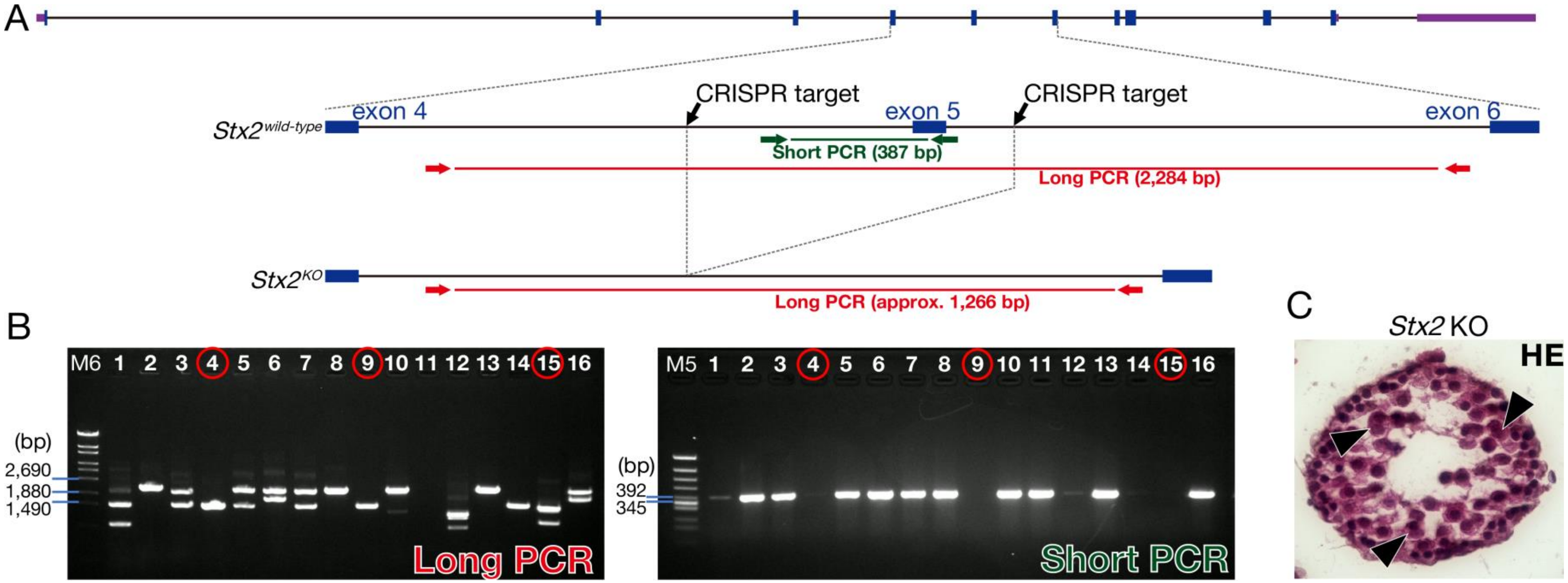


figure supplement 4. The production of the *Stx2* KO.

(A) *Stx2* KO mice generated by the CRISPR-Cas9 genome editing in mouse zygotes. Two CRISPR targets located 482 bp upstream and 148 bp downstream of exon 5 of *Stx2*. Arrows indicate long and short PCR primers for detecting the deletion of exon 5 of *Stx2*. (B) Electrophoresis data of PCR genotyping. We used #4, 9, and 15 as the Stx2 KO mice. M5: Marker 5. M6: Marker 6. (C) H&E staining of *Stx2* KO testis. Consistent with reported *Stx2* null phenotype (Fujiwara et al., 2013), our *Stx2* KO also showed aggregates of syncytia (AGS) (arrowheads).

**References**

Fujiwara, Y., Ogonuki, N., Inoue, K., Ogura, A., Handel, M. A., Noguchi, J., & Kunieda, T. (2013). t-SNARE Syntaxin2 (STX2) is implicated in intracellular transport of sulfoglycolipids during meiotic prophase in mouse spermatogenesis. *Biol Reprod, 88*(6), 141. doi:10.1095/biolreprod.112.107110

Muranishi, Y., Terada, K., Inoue, T., Katoh, K., Tsujii, T., Sanuki, R., . . . Furukawa, T. (2011). An essential role for RAX homeoprotein and NOTCH-HES signaling in Otx2 expression in embryonic retinal photoreceptor cell fate determination. *J Neurosci*, *31*(46), 16792-16807. doi:10.1523/JNEUROSCI.3109-11.2011
